# Standardized Informatics Computing Platform for Advancing Biomedical Discovery Through Data Sharing

**DOI:** 10.1101/259465

**Authors:** Vivek Navale, Michelle Ji, Evan McCreedy, Tsega Gebremichael, Alison Garcia, Leonie Misquitta, Ching-Heng Lin, Yang Fann, Matthew McAuliffe

## Abstract

**Objective:** The goal is to develop a standardized informatics computing system that can support end-to-end research data lifecycle management for biomedical research applications.

**Materials and Methods:** Design and implementation of biomedical research informatics computing system (BRICS) is demonstrated. The system architecture is modular in design with several integrated tools: global unique identifier, validation, upload, download and query tools that support user friendly informatics system capability.

**Results:** BRICS instances were deployed to support research for improvements in diagnosis of traumatic brain injury, biomarker discovery for Parkinson’s Disease, the National Ophthalmic Disease Genotyping and Phenotyping network, the informatics core for the Center for Neuroscience and Regenerative Medicine, the Common Data Repository for Nursing Science, Global Rare Diseases Patient Registry, and National Institute of Neurological Disorders and Stroke Clinical Informatics system for trials and research.

**Discussion:** Data deidentification is conducted by using global unique identifier methodology. No personally identifiable information exists on the BRICS supported repositories. The Data Dictionary provides defined Common Data Elements and Unique Data Elements, specific to each of the BRICS instance that enables Query Tool to search through research data. All instances are supported by the Medical Imaging Processing, statistical analysis R, and Visualization software program.

**Conclusion:** The BRICS core modules can be easily adapted for various biomedical research needs thereby reducing cost in developing new instances for additional biomedical research needs. It provides user friendly tools for researchers to query and aggregate genetic, phenotypic, clinical and medical imaging data. Data sets are findable, accessible and reusable for researchers to foster new research on various diseases.

## BACKGROUND AND SIGNIFICANCE

Biomedical research (basic, translational and clinical) requires an integrated systems approach for collecting, managing, and analyzing heterogeneous data. This need is especially critical in the translational research domain, where integration of large scale, multi-dimensional clinical phenotype and bio-molecular data sets can address fundamental research questions. Therefore, developing knowledge-based systems with a standard data dictionary, common data elements with domain repositories are needed to assist in tasks that require data reproducibility, scalability and accessibility for immediate and extended research.[1]

Various electronic data capture tools have been designed for supporting clinical and translational research. For example, the Research electronic data capture (REDCap) method is widely used to capture electronic medical data, [2] and information about the advantages and limitations of REDCap is available here.[3]

For clinical trial data management, the ‘OpenClinica’ provides a web-based electronic Case Report Form tool that can be accessed from most locations in the world.[4] However, OpenClinica does not support advanced data processing, validations, quality control functionalities that are needed as part of a biomedical information management system.[5]

The National Institutes of Health (NIH) is also committed to making public access to digital scientific data the standard for all NIH funded research.[6]

Knowledge discovery and innovation can be greatly facilitated when data is findable (F), accessible (A), interoperable (I) and reusable (R). The FAIR principles serve as a guide for data producers, stewards and disseminators for enhancing reusability of data, and are inclusive of data algorithms, tools and workflows that are essential for good data life cycle management.[7] Most importantly, sharing data can accelerate research by allowing re-analysis, aggregation, and rigorous comparison with other data, tools, and methods.

Prior to development of the Biomedical Research Informatics Computing System (BRICS) platform researchers engaged in biomedical fields such as the traumatic brain injury used disparate systems with different data elements that made data access and sharing difficult. To address some of these challenges, developing informatics systems with common data element (CDE) definitions, standards, and user-friendly interfaces for uploading, accessing and analyzing data became necessary, concepts involving unique subject identifier, data elements and data dictionary were developed and tested for the National Database for Autism Research (NDAR system).[8] The success of these concepts led to design and development of an integrated BRICS platform that can be reused and deployed for a wide range of biomedical research programs.

The BRICS platform works to support research networks that help biomedical researchers collect, validate, categorize, and analyze research datasets on a worldwide basis. The platform helps ensure there are fewer isolated, disparate research datasets as well as allows researchers to aggregate and query genetic, phenotypic, clinical, and medical imaging data.[9]

In addition, BRICS is intended to support the NIH Big Data to Knowledge (BD2K) initiatives such as the development of biomedical and healthCAre Data Discovery Index Ecosystem (bioCADDIE), aimed to enhance discoverability of biomedical datasets across multiple repositories.[10]

### OBJECTIVE

This paper discusses development of BRICS, a knowledge based informatics system with characteristics that include – (a) supporting domain specific metadata collection, (b) ability for assignment and retrieval of data by unique data identifiers, (c) capability to maintain data provenance (d) enabling standardized communications protocol, and (e) implementing community established standards for making data FAIR.

Making data FAIR fosters – (a) testing of new hypotheses, (b) allowing multi-study data aggregation, and (c) identify patterns not easily extracted from a single study.

Also, central to the objective of this work is to demonstrate implementation of disease-agnostic system capability with modular design to allow for use of one or all the BRICS functionality for various biomedical research program requirements.

## MATERIALS AND METHODS

### BRICS platform and architecture

The BRICS system is a collaborative and extensible web-based system to aid and support the collection of research studies and clinical trials. It consists of a collection of modular components which include: Account Management, the Global Unique Identifier (GUID) tool, the Data Dictionary Tool (DDT), Data Repository, Meta Study, Protocol and Form Research Management System (ProFoRMS) tool, Medical Image Processing, Analysis, and Visualization (MIPAV tool), and the Query Tool.

To ensure data security, the system is designed to employ multiple tiers of data security based on the content and level of risk associated with the data. The BRICS platform is Federal Information Security Modernization Act (FISMA) Moderate level compliant. The platform maintains confidentiality of research subjects but does provides information on the data and study protocols, to promote scientific collaboration by use/reuse of data. The various modules that comprise the BRICS architecture are shown in Figure 1 and brief description is provided below -

**Figure 1.**
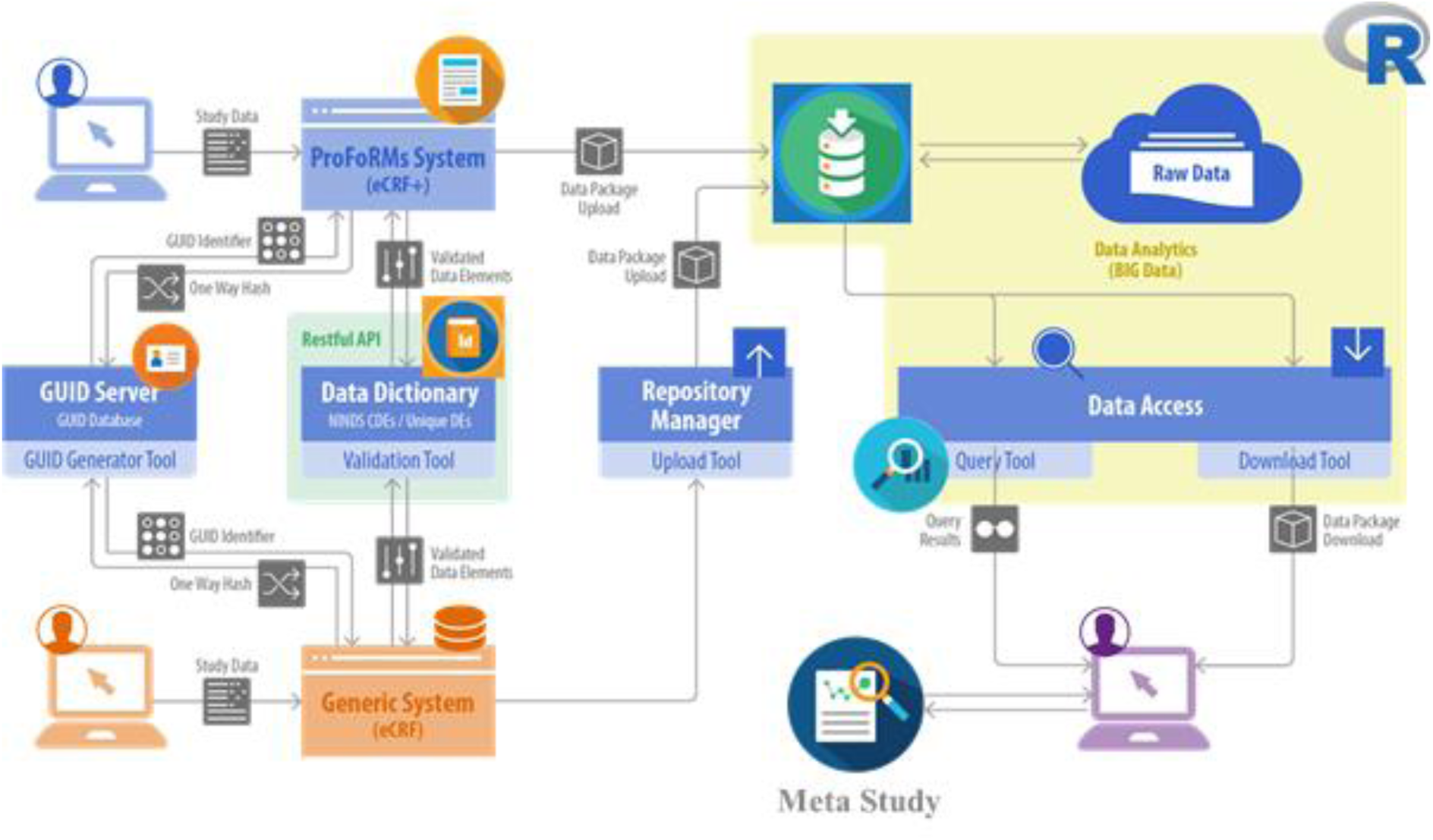
– BRICS Platform Architecture

#### Global Unique Identifier

Prior to data submission to the BRICS instance, data is deidentified by applying a unique code known as Global Unique Identifier (GUID).[11] The GUID is a computer-generated random alphanumeric code unique to each research participant (i.e., each person’s information has a different GUID). The GUID is generated at the researcher’s site, by collecting data using eForm and/or ProFoRMs, applying the GUID generator tool that is available as client software program (shown in Figure 1). The GUID database for each of the BRICS instances (currently seven) is maintained separately on GUID servers located at NIH.

By using of the GUID tool, no personally identifiable information is sent to the data repository, which minimizes identification risks to study participants, by keeping an individual’s information separate from that of another person and does not use names, addresses, or other identifying information. The unique code also allows BRICS to link together all submitted information on a single participant, giving researchers access to information that may have been collected elsewhere.

#### ProFoRMS

allows users to manage protocols by entering and editing protocol study information, attach necessary files, use the GUID for subject ID’s, support single site or multi-site protocols. It also creates and manages patient visit types. ProFoRMS interacts closely with the Data Dictionary, to facilitate researchers needs, to access non-copyrighted eForm’s for use. Other capabilities include scheduling for subject visits, collecting data, and eForm’s to add new data or modify previously collected data entries. It also allows for data to be viewed and discrepancies (e.g. double data entry) to be resolved, including all changes are tracked in the system that can be viewed in audit logs.

Users can also import data to BRICS repository by using their generic system data capture forms, after validation with the Data dictionary (as shown in Figure 1).

#### Data Repository

The BRICS Data Repository is the central hub of the BRICS system, providing functionality for defining and managing study information as well as for contributing, uploading, and storing the research data associated to each study (Figure 1). When an investigator is authorized to submit data to a BRICS instance, they can organize one or many datasets into a single entity called a Study. In general terms, a ‘Study’ is a container for the data to be submitted, allowing an investigator to describe, in detail, any data collected, and the methods used to collect the data. In addition, the repository module provides download statistics for specific studies, enabling research investigator to obtain information on their respective data that has been downloaded for other research activities, and overall increase data sharing and collaboration for additional research goals.

#### Data Dictionary

At the core of BRICS is data standardization approach, which makes it easier to compile and search large databases of research from diverse fields. The data dictionary provides defined Common Data Elements (CDEs) as well as Unique Data Elements (UDEs) specific to a given instantiation of the BRICS system, as well as translation across terms. For example, the Federal Interagency Traumatic Brain Injury Research (FITBIR) data dictionary incorporates and extends the CDE definitions developed by the National Institute of Neurological Disorders and Stroke (NINDS) CDE Project.[12] The CDEs help to make data findable that is most relevant to researcher’s needs, as well as promote reusability.

The data dictionary is comprised of three key components: data elements, form structures and eForms. A data element has a name, precise definition, and clear permissible values (codes), if applicable. A data element directly relates to a question on a paper/electronic Form (eForm) or field(s) in a database record. Form structures are the container to data elements and common data elements and correspond to a paper form. The form structures are later used to create eForm’s, which can be used in ProFoRMS based protocols for submitting research data via web portal.

Creating and editing Data Elements is part of the Data Dictionary tool, which enables browsing, filtering, auditing and searching for Unique Data Elements and Common Data Elements, and export Data Element capabilities. The Form Structure functionality enables to create, edit, browse, filter, search, import, export, publish, and archive various Form Structures (FS). Importantly data elements can be exported to REDCap format.

#### Query Tool

The Query Tool enables users to browse studies and forms, to select clinical data, and to search data within forms using a rich set of features to filter, sort, and combine records. Using GUID and a standard vocabulary via CDEs in forms, the Query Tool provides an efficient means to search through volumes of aggregated research data across studies to find the right datasets, to download and perform offline analysis using other tools (e.g. R, SAS, SPSS, etc.). The statistical ‘R-box’ has been incorporated in the Query Tool, to support analysis without having to download data.

The query tool has many ways to search for data. By default, the user is presented with all studies in the data repository that have data submitted against them. Users can search the query tool for desired data by searching by study, or across studies by form or an individual data element (Figure 2a).

**Figure 2a.**
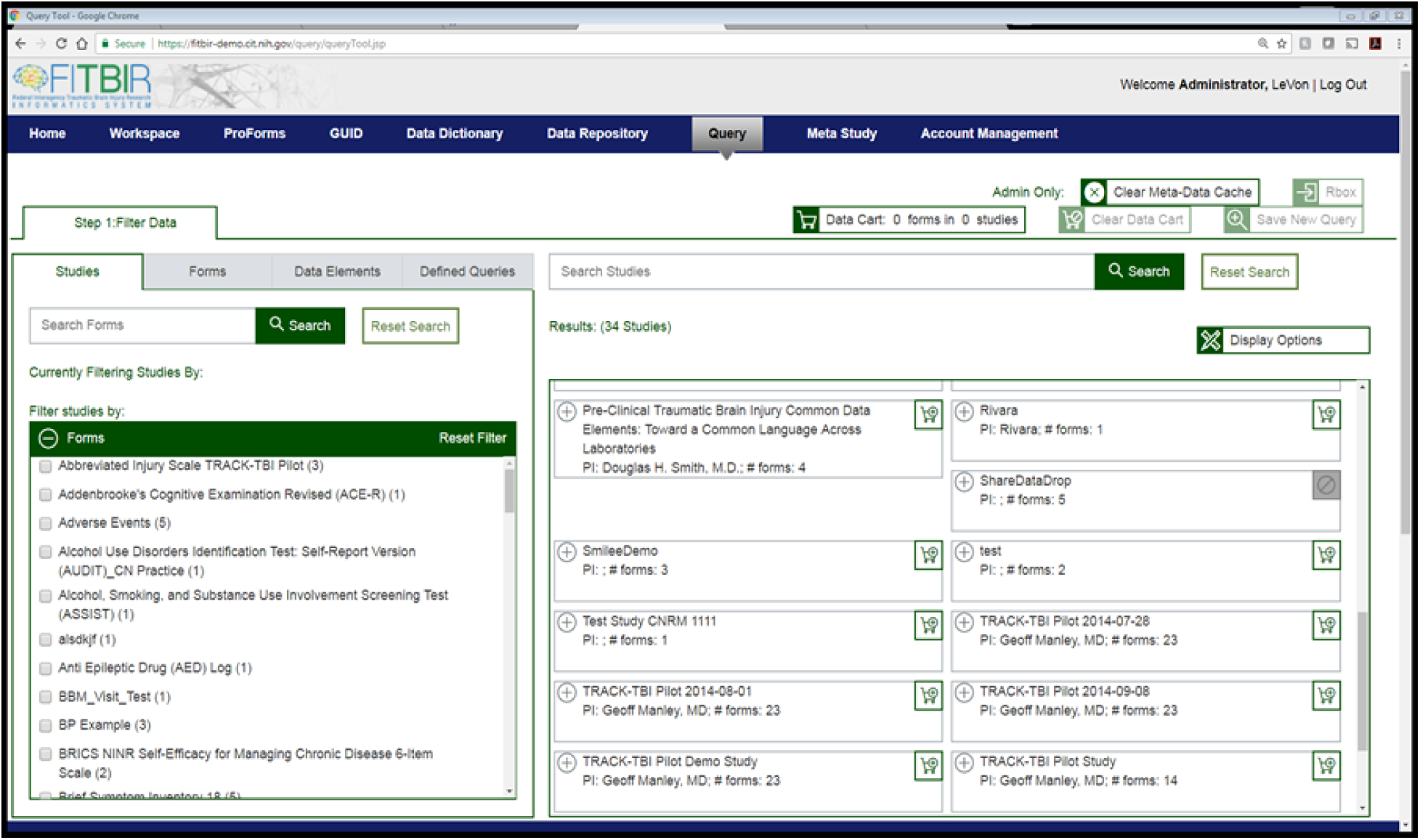
The Query Tool functionality is used to browse studies and forms, search data within forms and across studies (shown here for FITBIR).

Each column of data in a Query Tool result represents a well-defined element in the Data Dictionary, users can refine results by selecting from the list of allowed element permissible values, like male or female, or move sliders to select a range of numeric values, like age or outcome scores (shown in Figure 2b).

**Figure 2b.**
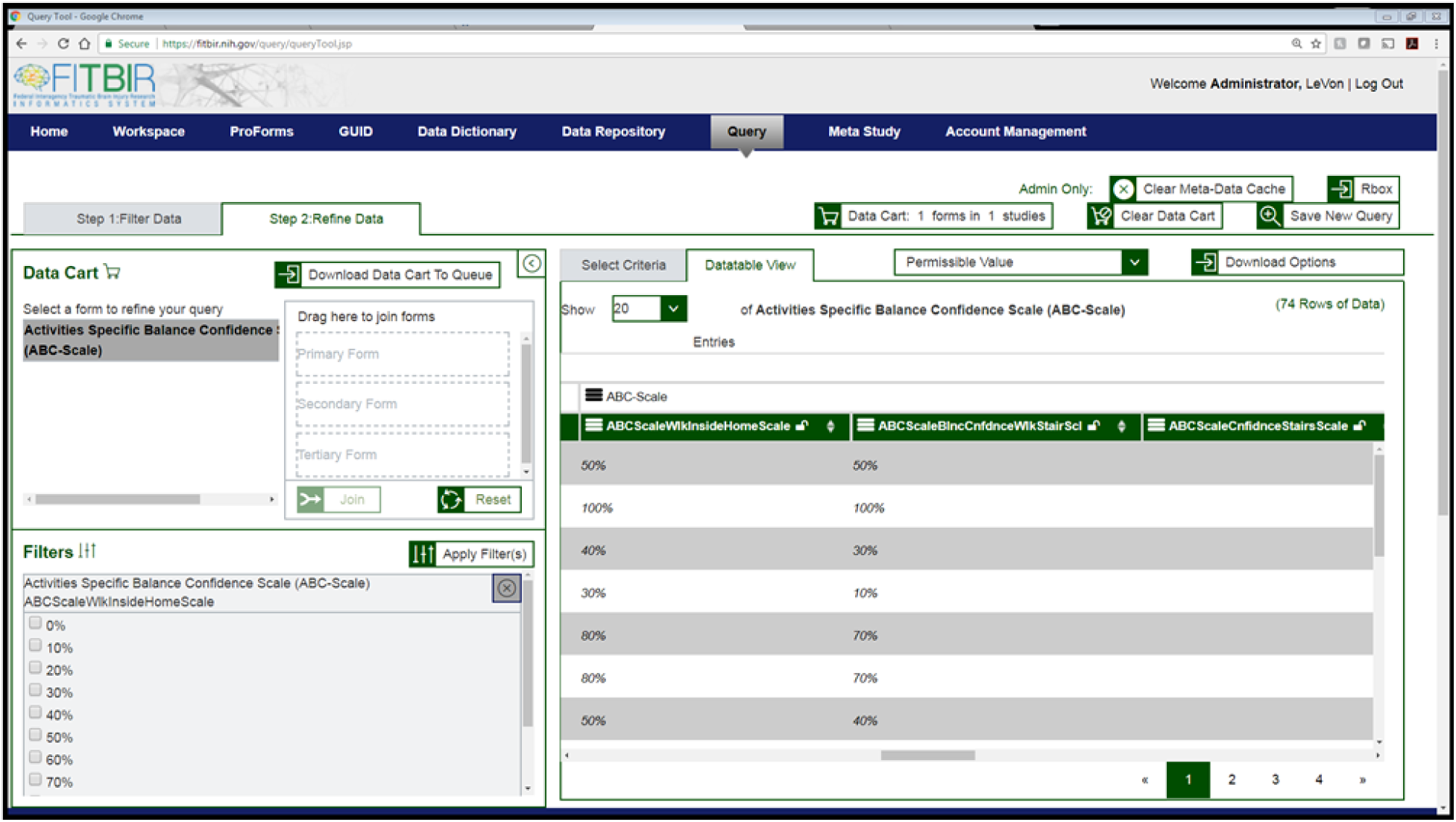
– The Query Tool can be used by users to select from a list of data elements that exist or are part of form structure.

In addition to providing tools to aid data discovery, the Query tool supports interactive features that facilitate analysis and practical use of the data through layered capability, based on the data element type. Varied data sets such as clinical, cognitive, demographic, and other phenotypic types are available in a BRICS instance and can be shared in CSV file format for download, storage in the Meta Study module of BRICS for further research and analysis.

BRICS uses Semantic Web technologies, to enable users to find, share, and combine information more easily. The ability to search by facets and join study data by subject IDs provides unparalleled capability for researchers to query data from the system without having to open an additional analytical tool.

#### Meta-Study

Findings from multiple other studies can be aggregated as a “Meta-Study” by researchers to conduct additional analysis in support of making data FAIR. Once a Meta-Study is published, a digital object identifier (DOI) is generated and associated with the Meta-Study. The information and more importantly the data within the Meta-Study can be referenced in publications. Current efforts are underway to make the DOI’s for Meta-Study be searchable via bioCADDIE.

#### MIPAV

is open source software image analysis application that can be used on any Java-compatible platform, including Windows, Apple OS X, and Linux. It supports over thirty file formats commonly used in medical imaging, including DICOM and NIfTI, and more than ten 3D surface mesh formats.[13] It also supports multi-scale and multi-dimensional image research from microscopy, to CT, PET, and MRI. BRICS’s inclusion of the MIPAV tool gives powerful capabilities for uploading image packages, as well as image analysis, not found in other informatics systems.[14]

The BRICS Image Submission Package Creation Tool leverages the medical image file readers in MIPAV to enable the mapping of image header data onto the data elements in imaging form structures for submission to the Data Repository.

## RESULTS

BRICS was initially deployed to support the U.S. Department of Defense’s (DoD) and the National Institute of Neurological Disorders and Strokes (NINDS), Federal Interagency Traumatic Brain Injury Research (FITBIR) project, and later for Parkinson’s Disease Biomarker Program (PDBP), the National Eye Institute (NEI) eyeGENE program, National Institutes of Nursing Research (NINR), Common Data Repository for Nursing Science (cdRNS), and three other smaller projects. Each data resource is separately branded but reuses the BRICS components, significantly cutting costs, and reducing development time. The knowledge based informatics system approach enables BRICS improvements to be quickly propagated to the various instances.

Currently there are seven BRICS instances for various biomedical research programs (Table 1 provides functionalities in use).

**Table 1.**
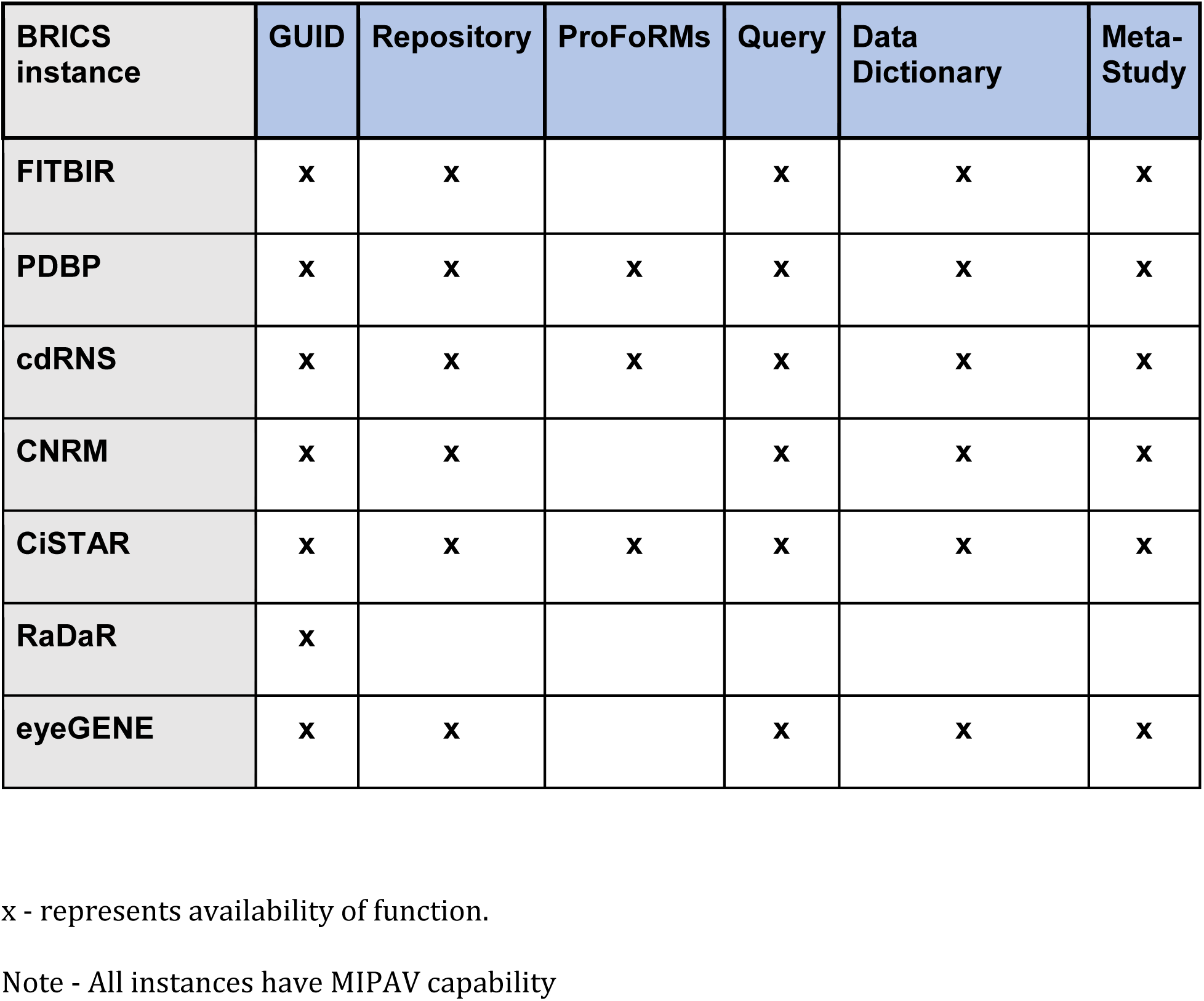
–Mapping of the BRICS instances with functionalities.

A brief description of the programs where BRICS have been deployed are provided below -

**Federal Interagency Traumatic Brain Injury Research (FITBIR) –** is a BRICS instance developed to advance comparative effectiveness research in support of improved diagnosis and treatment for those who have sustained a traumatic brain injury (TBI).[15] FITBIR stores data provided by TBI researchers, can accept high quality research data from more than 150 studies, regardless of funding source and location. The Department of Defense (DOD) and NINDS provided funding for TBI human subject studies (both retrospective and prospective) and have required their grantees to upload their clinical, imaging, and genomic data to FITBIR. As of 2017, there are 157 studies in FITBIR, spanning nearly hundred Principal Investigators (PIs), dozens of universities and research systems, the DOD, the National Institutes of Health. Data on 60,208 subjects, including 31,389 clinical image data sets are 3D volumes that are part of the repository. Currently there are a total of 1,857,926 records in FITBIR. Data provided to FITBIR for broad research access are expected to be made available to all users within six months after the award period ends.

**Parkinson’s Disease Biomarkers Program Data Management Resource (PDBP DMR)** -is a BRICS instance developed to support new and existing research and resource for promoting biomarker discovery for Parkinson’s disease. At the center of the PDBP effort is its Data Management Resource (DMR). The PDBP DMR uses a system of standardized data elements and definitions,[16] which makes it easy for researchers to compare data to previous studies, access images and other information, and order bio samples for their own research. PDBP’s needs have accelerated BRICS system development, such as enhancements to the ProFoRMS data capture module, also with an investment into a BRICS plug-in for managing bio samples. The PDBP DMR now contains over 1,500 enrolled subjects, 1,415 of whom have biorepository samples. Also, PDMP has currently a total of 55, 400 records.

**eyeGENE –** has a BRICS instance to support the National Ophthalmic Disease Genotyping and Phenotyping Network.[17] It is a research venture created by the National Eye Institute (NEI) to advance studies of eye diseases and their genetic causes, by giving researchers access to DNA samples and clinical information. Data stored in eyeGENE is cross-mapped to Logical Observation Identifiers Names and Codes terminology (LOINC) interoperability data standards.[18] Currently, eyeGene has 146, 024 records with 6,400 enrolled subjects.

**Informatics Core of Center for Neuroscience and Regenerative Medicine (CNRM) –** has a BRICS instance to support the CNRM medical research program with collaborative interactions between the U.S. DoD, NIH, and the Walter Reed National Military Medical Center. The Informatics Core provides services such as electronic data capture and reporting for clinical protocols, participation in national TBI research and data repository community, integration of CNRM technology requirements, and maintenance of a CNRM central data repository.[19] In addition, the Informatics Core has played an important role of providing technical and clinical expertise in steering the development of multiple BRICS modules used by FITBIR.

**Common Data Repository for Nursing Science (cdRNS) –** has a BRICS instance to support the NINR mission – to promote and improve the health of individuals, families, and communities.[20] To achieve this mission, NINR supports and conducts clinical and basic research and research training on health and illness, research that spans and integrates the behavioral and biological sciences, and that develops the scientific basis for clinical practice. [21] The NINR is a leading supporter of clinical studies in symptom science and self-management research. To harmonize data collected from clinical studies, NINR is spearheading an effort to develop CDEs in nursing science. Currently, there are 1358 records in cdRNS instance of BRICS.

**The Rare Diseases Registry –** has a BRICS instance for the Rare Diseases Registry (RaDaR) program of the National Center for Advancing Translational Sciences (NCATS). It is designed to advance research for rare diseases.[22] Because many rare diseases share biological pathways, analyses across diseases can speed the development of new therapeutics. The goal is to build a Web-based resource that integrates, secures and stores de-identified patient information from many different registries for rare diseases, all in one place.

**Clinical Informatics System for Trials and Research (CiSTAR) –** CiSTAR is a NIH intramural collaborative development project for a clinical research information system at NINDS. Using five BRICS modules, CiSTAR has merged two existing systems into one to support clinical trials and research.[23]

## DISCUSSION

The BRICS platform is comprised of 7 modules (shown in Figure 3) and are configured to form the full BRICS architecture, described in A major strength of the BRICS platform is that modules are disease-agnostic and reusable for various applications, a feature that translates to cost efficiency, by modules being relatively quickly configured and deployed to meet specific needs of new research project. These modules can be deployed as one integrated system, or as a subset of components.

**Figure 3.**
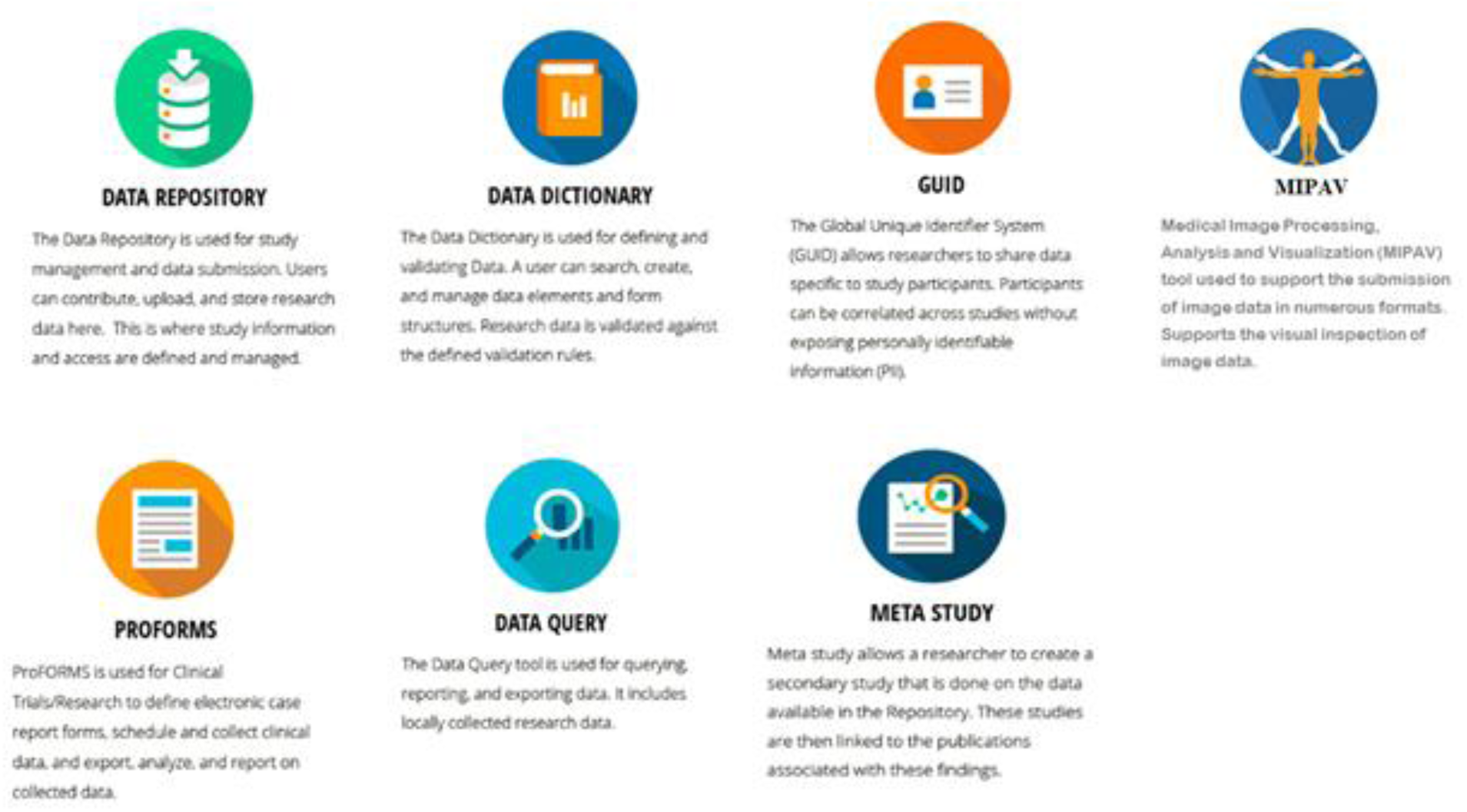
– Major BRICS Modules

Easy access to the modules in BRICS is provided by respective icons -

BRICS system changes and enhancements can be rapidly propagated across all the various instances to facilitate protocol initiation, data collection, data management and data analysis. The ProFoRMS tool and Data Dictionary have proven to save time and enhance data standardization. Reusability has enabled substantial cost reductions for system development with flexibility to develop and deploy multiple biomedical research data repositories.

BRICS also supports clinical data submissions formats of tab delimited or comma separated values (CSV). All data (clinical assessment, imaging, and genomics) submitted to a BRICS instance, whether through ProFoRMS, MIPAV, or via semi-automated method is transferred as a CSV file format, that must be validated against the Data Dictionary prior to upload. This validation step serves as a form of data quality control. Submissions are validated against CDE data definitions (such as permissible values, data input restrictions, min/max values, etc.), and data cannot be uploaded unless all validation checks have been corrected.

The BRICS Validation Tool checks the data proposed for submission against the Data Dictionary to ensure that the data comply with the established standards. After using the validation tool to validate their data against the Data Dictionary and its CDEs, the users can upload data into their BRICS instance study folder that is in a private state. While in the private state, only approved users, who have been granted access by the owner of the study, have query access to the data. After a period defined by a particular BRICS instance policy, the data enter a new shared state. All approved users have query access to the data in the shared state

The modular architecture for BRICS provides users to select the functionalities needed for the instance. As shown in Table 1 BRICS functionalities can be deployed for various biomedical applications. PDBP, cdRNS, and CiSTAR use all the BRICS functionality, FITBIR, CNRM and eyeGENE use their generic system for data capture and utilize the rest of BRICS functionality. The RaDaR uses the GUID for their current system in use with their storage repository.

BRICS uses Resource Description Framework (RDF) as the standard model for data representation on the Web.[24] Developed under the guidance of the World Wide Web Consortium (W3C), RDF is an infrastructure that enables the encoding, exchange and reuse of structured metadata on the web. It provides a means for publishing both human-readable and machine-processable vocabularies designed to encourage the reuse and extension of metadata semantics among different information communities. RDF identifies any resource by an Uniform Resource Identifier (URI), and describes it with XML statements comprising a subject (resource), a predicate (property), and an object (property value, can be atomic or linked to another resource). The subject->predicate->object binary relationship is usually referred to as a “triple”. An RDF database is made up of a collection of linked triples, and so is named as RDF triplestore. Using this simple model, structured and semi-structured data can be integrated, exposed to users, and shared across different applications. Like relational databases, data stored in an RDF triplestore can be queried with RDF query language. BRICS uses SPARQL Protocol and RDF Query Language (SPARQL) to retrieve and manipulate semantic metadata, SPARQL has been a standard RDF query language and officially recommended by W3C since 2008.

Data studies in a BRICS instance also have citable digital object identifiers/uniform resource identifiers (DOIs/URIs) functionality. BRICS instance studies include detailed metadata about a study and provide the container for associated dataset submissions. The metadata is referenced by DOI, that is accessible via static URL, including references to datasets from contributing studies, to ensure that they receive appropriate attributions with capability to track their uses. The meta study module of the BRICS system uses the Data Tag Suite (DATS) specifications.[25]

Use of common data elements for each of the BRICS instances provides for data harmonization, collection and management are enabled by data Dictionary that facilitates in integration of data for addressing newer research questions.

## CONCLUSION

BRICS platform provides the basis for developing knowledge based informatics system that can be used for biomedical research applications. The basic architecture and core functionality is easily adaptable for various research needs thereby reducing development cost for newer instances. The standardized approach of using common data element and data dictionary for each of the instances promotes data harmonization, makes data sets to be findable, accessible and reusable for researchers to foster new research on various diseases. Additionally, it provides users with friendly tools for researchers to query and aggregate genetic, phenotypic, clinical and medical imaging data within a secure environment.

## Acknowledgments

The Research is supported by the Intramural Research Program, Center for Information Technology, the National Institutes of Health. The opinions expressed in the paper are those of the authors and do not necessarily reflect the opinions of the National Institutes of Health.

## No Conflict of Interest Declaration

The authors have no competing interests.

## Abbreviations/ Acronyms

BD2K: Big Data to Knowledge
bioCADDIE: The biomedical and healthCAre Data Discovery Index Ecosystem
BRICS: Biomedical Research Informatics Computing System
CDE: Common Data Element
cdRNS: Common Data Repository for Nursing Science
CiSTAR: Clinical Informatics System for Trials and Research
CNRM: Informatics Core of Center for Neuroscience and Regenerative Medicine
DATS: Data Tag Suite
DAQC: Data Access and Quality Committee
DDT: Data Dictionary tool
DOD: U.S. Department of Defense
DOI: Digital Object identifier
eForm: Electronic Form
eyeGene: The National Ophthalmic Disease Genotyping and Phenotyping Network
FAIR: Findable, Accessible, Interoperable, Reusable.
FDA: U.S. Food and Drug Administration
FISMA: Federal Information Security Modernization Act
FITBIR: Federal Interagency Traumatic Brain Injury Research
FS: Form Structures
GUID: Global Unique Identifier
MIPAV: Medical Image Processing Analysis and Visualization
NDAR: National Database for Autism Research
NEI: National Eye Institute
NIH: National Institutes of Health
NINR: National Institute of Nursing Research
NINDS: National Institute of Neurological Disorders and Stroke
PDBPDMR: Parkinson’s Disease Biomarker Program Data Management Resource
ProFoRMS: Protocol and Form Research Management System
PTSD: Posttraumatic stress disorder
RaDaR: Rare Diseases Registry Repository
REDCap: Research Electronic Data Capture
RDF: Resource Description Framework
TBI: Traumatic brain injury
UDE: Unique data element
URI: Uniform resource identifier

